# Pathological musculoskeletal characterisation of a new model of spontaneous OA in CD1 mice

**DOI:** 10.64898/2026.10.12.758449

**Authors:** Lorenzo Ramos-Mucci, Emily Shorter, Tayfun Dikmen, Craig Keenan, Yan Wen, Yanhong Wang, Eithne Comerford, Katarzyna Goljanek-Whysall, Blandine Poulet

## Abstract

Osteoarthritis (OA) is a whole-joint disease involving pathological changes in cartilage, synovium, ligaments, menisci and periarticular muscle. This study provides a comprehensive musculoskeletal characterisation of spontaneous OA in male CD1 mice to evaluate their suitability as a model of naturally occurring disease. Knee joints from 3-4 and 6-8 month-old mice were analysed histologically, OARSI scoring and immunohistochemistry for markers of chondrogenesis and endochondral ossification. Quadriceps and gastrocnemius muscles were examined for fibre cross-sectional area.

CD1 mice displayed mild OA at 3-4 months, progressing to moderate-to-severe disease by 6– 8 months. Cartilage loss, synovial hyperplasia and fibrosis increased across compartments. Anterior cruciate and collateral ligaments, and medial meniscus exhibited marked pathological remodelling, including proteoglycan-rich rounded-cell regions and evidence of endochondral ossification with increased COL2, SOX9, RUNX2 and ASPN expression. Gastrocnemius muscle showed significant atrophy at 6-8 months, correlating with OA severity, whereas quadriceps atrophy was not significant.

These findings establish CD1 mice as a robust spontaneous OA model that recapitulates the multi-tissue complexity of human disease. Because CD1 mice are globally available, cost-effective, genetically diverse and frequently, this model offers a practical and translationally relevant platform for investigating whole-joint organ mechanisms of OA progression and for evaluating therapies aimed at modifying disease across the entire joint.

## Introduction

Osteoarthritis (OA) is the most common chronic joint disorder and a leading cause of pain and disability worldwide, imposing substantial personal and socioeconomic burden (Man and Mologhianu, 2014). Despite its high prevalence, no disease-modifying OA drugs are currently available. Ageing is the strongest risk factor for OA, but obesity, joint injury and genetic predisposition also contribute significantly to disease onset and progression (Johnson and Hunter, 2014). OA is now recognised as a whole-joint disease rather than a condition confined to articular cartilage, involving coordinated pathological changes across cartilage, synovium, ligaments, menisci, subchondral bone and peri-articular muscles (Loeser et al., 2012). This multi-tissue complexity underscores the continued need for animal models that capture interactions between joint tissues during disease development.

Murine models are indispensable for dissecting OA pathogenesis and evaluating tissue-specific mechanisms. These models can be broadly classified into mechanical, ageing-related, spontaneous, obesity/metabolic and chemical induction models, reflecting known aetiologies of human OA (Kuyinu et al., 2016). Spontaneous models are particularly valuable because they mimic the natural, age-related progression of primary OA without surgical or chemical intervention. Ageing-associated OA arises gradually through intrinsic processes such as cartilage matrix deterioration, subchondral bone remodelling and low-grade inflammation (Loeser, 2010). Several mouse strains, including the Str/ort mouse, develop spontaneous OA with hallmark features such as osteophyte formation, cartilage erosion, subchondral sclerosis and synovial changes(Mason et al., 2001; Staines et al., 2017), making them valuable for investigating genetic and biomechanical contributors to disease.

Recent work has suggested that outbred CD1 mice develop spontaneous OA by approximately one year of age, exhibiting cartilage degeneration and osteophyte formation (A. Ratneswaran, 2018). However, pathological involvement of other joint tissues—including ligaments, synovium and peri-articular muscles—has not been characterised, and early disease stages remain undescribed. Given the increasing recognition that OA is a disease of the entire joint organ, models that capture multi-tissue pathology are essential for understanding cross-tissue signalling and whole-joint degeneration.

This study provides the first comprehensive characterisation of OA development in male CD1 mice. We define the temporal progression of articular cartilage degeneration and describe previously unreported pathological changes in synovium, knee ligaments, menisci andperi-articular skeletal muscles. These findings establish CD1 mice as a valuable spontaneous OA model that recapitulates the multi-tissue complexity of human disease and offers new opportunities for studying whole-joint mechanisms of OA pathogenesis.

## Materials and Methods

### Animals and Experimental Groups

All mice were kept in polypropylene cages of 2–6 littermates, subjected to 12-hour light/dark cycles at 21±2°C, and fed Rat and Mouse Breeder and Grower from Special Diets Services (Witham UK). All procedures complied with Animals (Scientific Procedures) Act 1986 and local ethics committee, including the University of Liverpool Animal Welfare and Ethical Review Body, under the Home Office Project Licences P267B91C3 and PP3119883. At the end of the experiments, mice were culled by cervical dislocation. Male outbred CD-1 mice (Charles River, UK) were used as a spontaneous model of OA and were culled at either 3-months (n=9), 4-months (n=5), 6-months (n=10), or 8-months (n=6). Ages were selected based on freely available cadaveric mouse samples within the University of Liverpool, Biological Services Unit, and to increase statistical power, we merged 3 and 4 months-old mice and 6 and 8 months old mouse samples for this study. This combining of age groups was supported by the OA scores showing no differences in disease severity between 3 and 4 months, nor between 6 and 8 months samples (Supplementary Figure 1).

### Knee joint histology

Knee joints were fixed in 10% neutral buffered formalin for 24 hours, washed in water and stored in 70% ethanol. For histological processing, joints were decalcified with 10% formic acid (Sigma) for 10 days, washed, dehydrated and processed for wax embedding. Knee joint samples were paraffin embedded coronally and sequentially sectioned at 6µm across the entire joint (Poulet et al., 2011).

Toluidine blue was used to assess pathophysiological changes of joint tissues, including articular cartilage (AC) loss, synovial pathology, ligament and meniscal structural pathology. A quarter of the sectioned histology slides across the joint were stained with toluidine blue at regular intervals (Poulet et al., 2011). Slides were dewaxed and hydrated, then pre-conditioned with sodium acetate buffer (0.1M, pH 5.6) and stained with toluidine blue (0.1% toluidine blue in 0.1M solution of acetate buffer, pH 5.6). Slides were then counterstained with 0.2% fast green, washed with distilled water, dehydrated with acetone and xylene.

Articular cartilage (AC) lesion severity was scored using the OARSI cartilage scoring system (Glasson et al., 2010) in each compartment (medial tibia (MT), medial femur (MF), lateral tibia (LT) and lateral femur (LF)). The average mean and maximum lesion scores were calculated for each compartment and summed scores of all the compartments were also calculated from the mean and maximum scores (Poulet et al., 2011).

Synovial pathology was scored according to the scoring system of (Obeidat et al., 2024) at medial tibial, medial femoral, lateral tibial and lateral femoral synovial gutters on the Toluidine Blue-stained sections. Synovium immediately bordering the meniscus or the bone surface was avoided during the assessments. Sections from the central region of the joint were examined and the most diseased section was selected for scoring. Each anatomical region was assessed for lining hyperplasia, subintimal cellularity and subintimal fibrosis. For each feature, scores from the four regions were summed to give a total joint score per sample, and each feature was analysed individually.

Ligaments (cruciate and collateral) and menisci were assessed for structural modifications, such as fibre alignment, proteoglycan staining (Toluidine blue) and cellular morphology arising from OA development, including cruciate and collateral ligaments(Ramos-Mucci et al., 2020).

### Immunohistochemistry

Immunohistochemistry was performed to localise expression of collagen type II (COL2), SRY-box transcription factor 9 (SOX9), Runt-related transcription factor 2 (RUNX2) and Asporin (ASPN), based on our published protocol (Ramos-Mucci et al., 2020). One to two adjacent slides per animal in at 3 months and 6 months of age (n=8 and n=10 respectively) were selected where cruciate and collateral ligaments were visible. Briefly, sections for COL2 immunostaining were preconditioned with 0.2M hydrochloric acid (15 minutes) and antigen retrieval applied with pepsin (3mg/mL in 0.02M hydrochloric acid; 45 minutes at 37°C). No antigen retrieval steps were performed for SOX9, RUNX2 and ASPN. All slides were washed with PBS-T (PBS with 0.1% Tween, Sigma) and blocked for endogenous peroxidase with 0.3% hydrogen peroxide (Sigma; 1h). After washing (PBS-T) endogenous Avidin/Biotin binding was blocked (Avidin/Biotin Blocking Kit; Vector Labs, SP2001). Non-specific binding sites were blocked with serum block of either mouse-on-mouse (MoM) blocking reagent (Vector Labs) for mouse primary antibodies (COL2) or 10% goat serum (Vector Labs) for rabbit primary antibodies (SOX9, RUNX2, ASPN; 1h). Primary antibodies were incubated overnight at 4°C and included: COL2 (1/100; Mouse monoclonal, Thermo, MA5-12789), SOX9 (1/1000; Rabbit polyclonal, Millipore, AB5535), RUNX2 (1/1000; Rabbit polyclonal, Abcam, ab23981), and ASPN (1/400; Rabbit polyclonal, Abcam, ab58741). Negative controls included a mouse IgG (2µg/mL, Sigma) or rabbit IgG (1µg/mL, Vector Labs; representative images can be found in Supplementary Figure 2). All antibodies were diluted in PBS. Following washing (PBS-T), a biotinylated secondary antibody (Vector Labs) was applied, followed by application of Vectastain (Vectalabs). Stains were developed with DAB Substrate Kit (Peroxidase) (Vector Labs), dehydrated and mounted with DPX mounting medium (Sigma).

### Skeletal muscles histology

For each mouse, gastrocnemius and quadriceps muscles from the right hindlimb were processed for histology: each muscle was halved laterally, mounted on an electron microscopy cork disc, embedded in Optimal Cutting Temperature compound, and cryoprotected in isopentane chilled with liquid nitrogen (Terry and Wells, 2016) . Sections were cut at 10 µm using a Leica CM1860 cryostat. For staining, sections were fixed using ice cold methanol, and stained with a 1:1000 dilution of rhodamine wheat germ agglutinin (WGA; 5 µg/mL; Vector Laboratories, UK) for 10 minutes. Vectashield HardSet Antifade Mounting Medium with DAPI (Vectorlabs, UK) was used to coverslip the section and protect against photobleaching.

For analysis of fiber cross-sectional area (CSA) in the gastrocnemius and quadriceps muscles, myovision software was used (Wen et al., 2018). CZA files of entire muscle sections were opened in Zen Blue (Zeiss) where then smaller images throughout the section were captured and saved as a TIFF file. A minimum of three images across the section were used for muscle fiber analysis. TIFF files were loaded into the Myovision software. Minimum and maximum area was set to 50 and 5000, respectively. The analysis was repeated for each image of the section and then data combined to obtain an average cross-sectional area across the section.

### Statistical analysis and normality of data testing

Normality and statistical analysis were calculated using GraphPad Prism (GraphPad v8). Normality of data was tested using a Shapiro-Wilk test (p-value ≤0.05 for statistical significance). When the data was normally distributed, an unpaired T-test was used, for not normally distributed data a Mann-Whitney test was used. Correlation between muscle fiber CSA and OARSI scores were calculated using Pearson correlation coefficient.

## Results

### CD1 mice exhibit significant spontaneous OA at 6-8 months-old compared to their younger counterparts

CD1 mice showed minimal OA at 3 and 4 months of age, and progressed to moderate and severe OA at 6-8 months old, with summed mean and maximum scores reaching significance (p<0.0001, Fig.1A and Fig.1C). All compartments were affected by cartilage degeneration, with more severe lesions in the medial tibia and least affected in the lateral femur (Fig.1B and Fig.1D). Cartilage lesions varied from maximum of 2 (mild lesions down to the intermediate zone of articular cartilage) in young mice and increased to the maximum score of 6 in a subset of mice in older mice (severe loss of cartilage with bone exposed). In line with OARSI scores of articular cartilage lesions, synovial pathologies were seen, including hyperplasia, cellularity and fibrosis (Fig.1I-J); all three showing significant increases between 3-4 months and 6-8 months joints (p=0.033, p=0.003, p<0.0001, respectively; Fig.1E-G) in both lateral and medial compartments. In addition, osteophytes developed on the margins of the joint, including fully ossified osteophytes in the severe joints (Fig.1I-J).

**Fig. 1:**
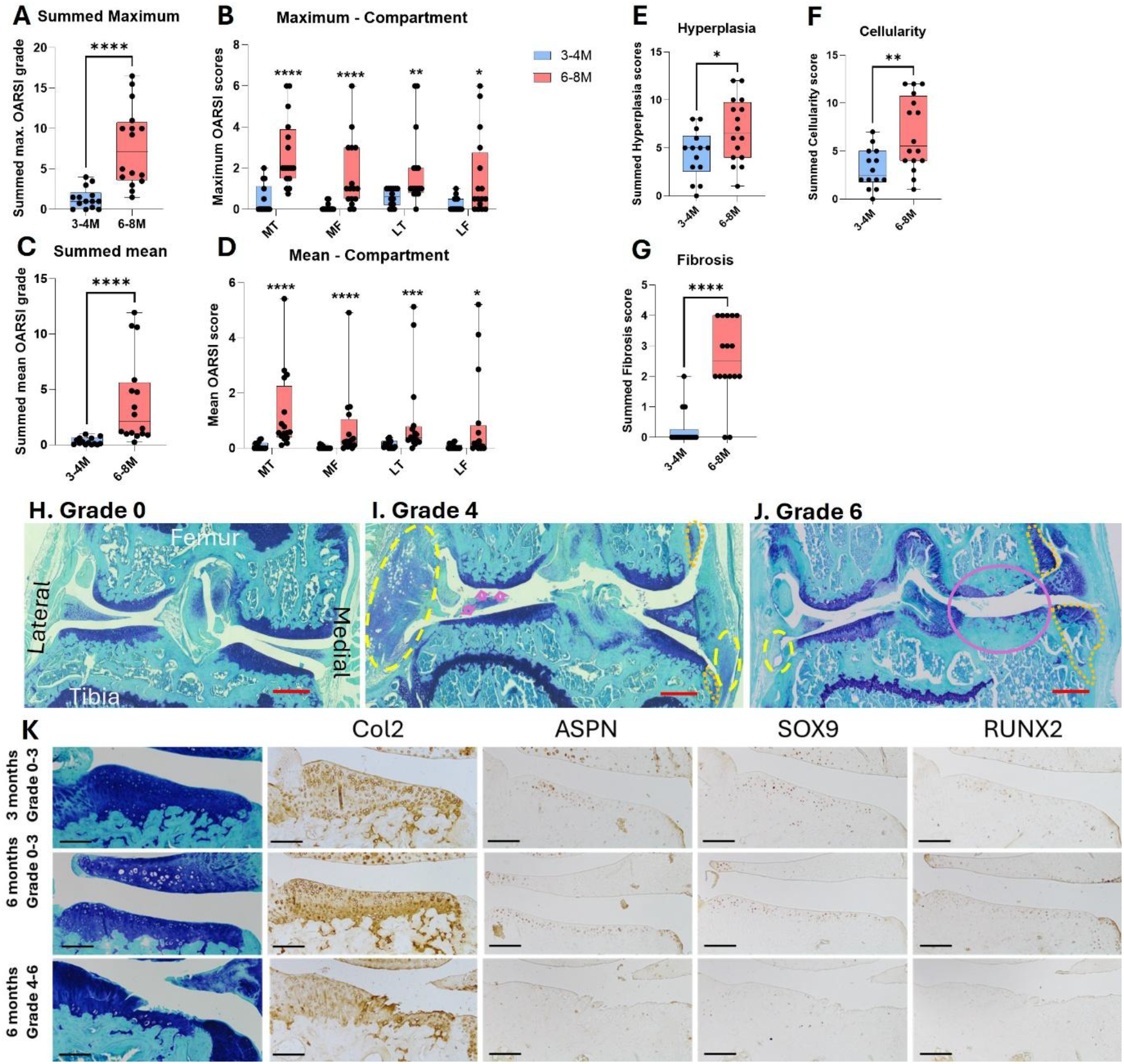
Histopathology of articular cartilage and synovium with OA development in CD1 mice. (A-D) OARSI scores of articular cartilage loss in measured as summed maximum (A) and summed mean(C) scores across all compartments, and separated compartments maximum (B) and mean (D) OARSI scores. (E-G) synovial pathology scores across all compartments of the knee, with hyperplasia (E), cellularity (F) and fibrosis (G) summed scores. (H-J) Representative histological coronal sections of CD1 knee joints stained with Toluidine Blue of typical OARSI grade 0 (H; healthy joint), grade 4 (I) moderate severity with cartilage loss (pink arrows), and grade 6 (J) severe OA joint with complete loss of AC on the medial femur and tibia (red circle) with underlying subchondral bone sclerosis. Osteophytes are delineated with orange dots on the medial femur and tibia, with early osteophytes (I) and large ossified osteophyte (J). Synovial hyperplasia is circled in yellow dashes on the lateral side of the joint showing hyperplasia and fibrosis. (K) Representative histological sections of the AC stained with Toluidine Blue, and immunolabelled for Col2, ASPN, SOX9 and RUNX2, in healthy AC at 3 (top row) and 6 months (middle row), and in OA sections at 6 months (bottom row). Scale bars = 50µm, black bars = 25µm.

Articular cartilage of CD1 mice showed expression of collagen type 2 across the whole cartilage, as well as cellular expression of SOX9, RUNX2 and ASPN (Fig.1K). All were similarly expressed in mild grade 2 cartilage but were lost from the whole condyle in more severe OA cartilage except for collagen type 2.

### Pathological endochondral ossification in the cruciate and collateral ligaments of osteoarthritic CD1 mice

Toluidine blue staining was used to analyse anterior cruciate ligament (ACL) and collateral ligaments structure and cellular morphology in 3-4 months-old and 6-8 months-old CD1 knee joints. Three-four months-old CD1 murine knee joints had an organised ACL structure with aligned collagen fibres and toluidine blue staining extending from the attachment site to the mid-ligament region (Fig.2A). The tibial attachment site had a delineated fibrocartilaginous zone with rounded-shaped cells, and spindle-shaped cells in the mid-ligament region (Fig.2A). In the 6-8 months-old CD1 knee joints with OA grade 4, ACL structure was modified with misalignment of the fibres and an increase in the ‘rounded’ cell types near the tibial attachment site and in the mid-ligament region (Fig.2B). In OA grade 6 CD1 knee joints, the ACL had complete toluidine blue staining in all regions of the ligament, suggestive of substantial proteoglycan deposition, and ‘rounded’ cell types near the tibial attachment site (Fig.2C). Immunostaining showed COL2 restricted to some areas of the ligament, similar to Toluidine blue stained proteoglycan-rich areas, with a relative absence of SOX9, RUNX2 and ASPN (Fig.2D). In contrast, OA ligaments (from joints with OARSI scores>3; Fig.2E) showed significant COL2 deposition throughout the ligament, with an increase in the rounded cells from the enthesis region deeper into the ligament, that also showed expression of endochondral ossification markers SOX9, RUNX2 and ASPN.

**Fig. 2:**
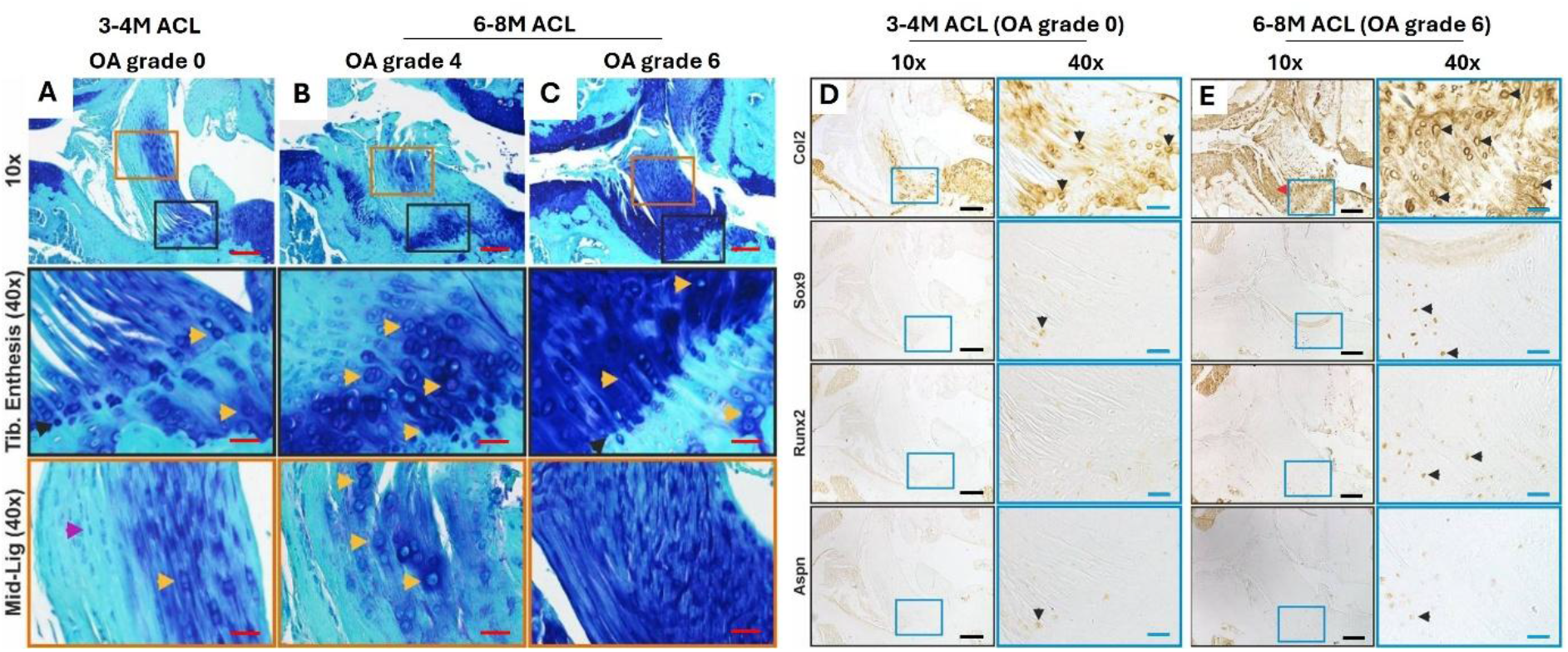
Immunostaining of the anterior cruciate ligament (ACL) tibial enthesis and mid-ligament in CD1 knee joints. Representative images of (A-C) Toluidine Blue stained sections of the whole ACL, and higher magnification of the tibial enthesis and mid-ligament areas in (A) a healthy 3 months-old CD1 (OARSI grade 0) and 6 months-old moderate (B; OARSI grade 4) and severe (C; OARSI grade 6) knee joint ACL. Yellow arrows point to rounded cells, red arrow points to spindle shaped fibroblasts. (D-E) Representative images of immunolabelled ACL in a healthy (D; OARSI grade 0) and OA joint (E; OARSI grade 6), for matrix staining with COL2, and cellular staining with SOX9, ASPN and RUNX2. Black arrows points to examples of positively immunostained cells. Scale is 100 µm (10x) and 25 µm (40x).

Similarly, healthy collateral ligaments in 3-4 months-old CD1 (grade 0) ligaments showed organised collagen fibre alignment and some toluidine blue staining at the tibial enthesis (Fig.3A). Meanwhile, collateral ligaments from OA joints (6-8 months-old) showed evidence of hyperplasia, toluidine blue staining (proteoglycan deposition) with the presence of rounded cells instead of spindle-shaped fibroblasts, and potential ossification in severely diseased joints (Fig. 3B-E). Enthesis went from highly organised with a clear tidemark demarcation of calcified and non-calcified areas, and fibre alignment along the ligament, to loss of clear tidemark boundaries at the ligament enthesis and of fibre alignment with increasing OA severities (Fig. 3B-E). Immunostaining of healthy 3-4 months-old CD1 knee joints demonstrated a collateral ligament femoral enthesis with organised COL2 staining and sparse expression of SOX9, RUNX2 and ASPN (Fig.3F). Absence of COL2, SOX9, RUNX2 or ASPN staining was also noted in the healthy mid-ligament region. Collateral ligaments in OA knee joints in severe OA (from grade 4 until 6 in 6-8 months-old) showed increased expression of these endochondral ossification markers at the femoral enthesis and in the mid-ligament region, especially in cells with ‘rounded’ morphology (Fig.3G).

**Fig. 3:**
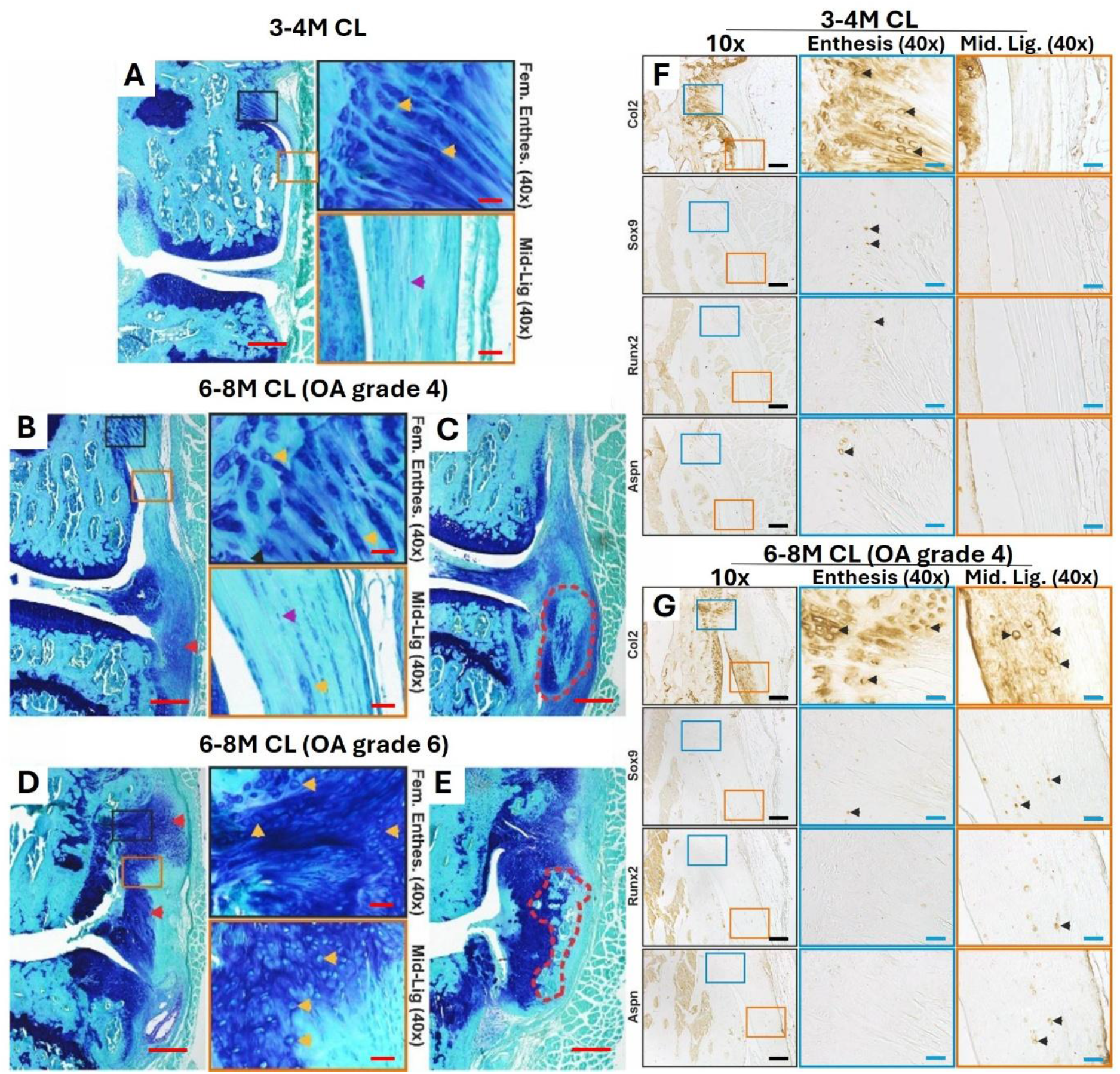
Histological toluidine blue and immune-staining of the collateral ligament (CL) in healthy and OA knee joints of CD1 mice. Representative images of toluidine blue stained sections (A-E) in:A) Healthy 3 months-old CD1 knee joints with organised and stained fibres (yellow arrows) near the femoral enthesis and spindle-shaped fibroblasts in the mid-ligament region (purple arrow). (B-C) OA grade 4 CD1 CLs with round cells in the mid-ligament region (yellow arrow) adjacent to areas with spindle-shaped fibroblasts (purple arrow), and areas of toluidine blue staining (red arrow) near the tibial attachment, that can become more severe with areas of chondroplasia (C; circled in red dashed lines). (D-E) OA grade 6 CD1 CLs with severe hyperplasia throughout the ligament, rounded cells (yellow arrows) and areas of strong toluidine blue staining (red arrows). Severely affected ligaments show areas of ossification (circled with red dashed lines). (F-G) Immunostaining of the CL for COL2, SOX9, RUNX2 and ASPN, with higher magnifications of the enthesis and mid-ligament regions, in (F) healthy 3 months-old CD1 knee and (G) in 6 months-old OA grade 4 CD1 knee. Black arrows point to examples of positive cellular staining. Scale is 200µm (4x, red bars), 100 µm (10x, black bars), 25µm (40x, blue bars and red).

### Osteoarthritic CD1 mice display pathological meniscal endochondral ossification

Healthy menisci in 3-4 months-old CD1 knee joints are organised in an outer hyaline cartilage layers, central ossified region and fibrous attachment (Fig.4A). At OA grade 4 and 6 of the 6-8 months-oldCD1 knee joints, menisci showed hyperplasia of the fibrous region with toluidine blue staining and ‘rounded’ cell morphology, as well as osteophyte formation at the anterior horn of the medial meniscus (Fig.4B-C). Immunostaining of the healthy 3-4 months-old CD1 menisci showed COL2 staining in the hyaline cartilage region and sparse COL2, SOX9, RUNX2, and ASPN staining in the fibrous region (Fig. 4D). In the enlarged fibrous region of the OA grade 4 CD1 medial menisci, COL2, SOX9, RUNX2 and ASPN staining were more prominent or evident in the outer fibrous regions and corresponding to areas with ‘rounded’ cell morphology (Fig.4E). As OA progressed (grade 6), COL2 deposition spread throughout the entire meniscus and SOX9, RUNX2 and ASPN were also present in the outer fibrous regions (Fig.4F).

**Fig. 4:**
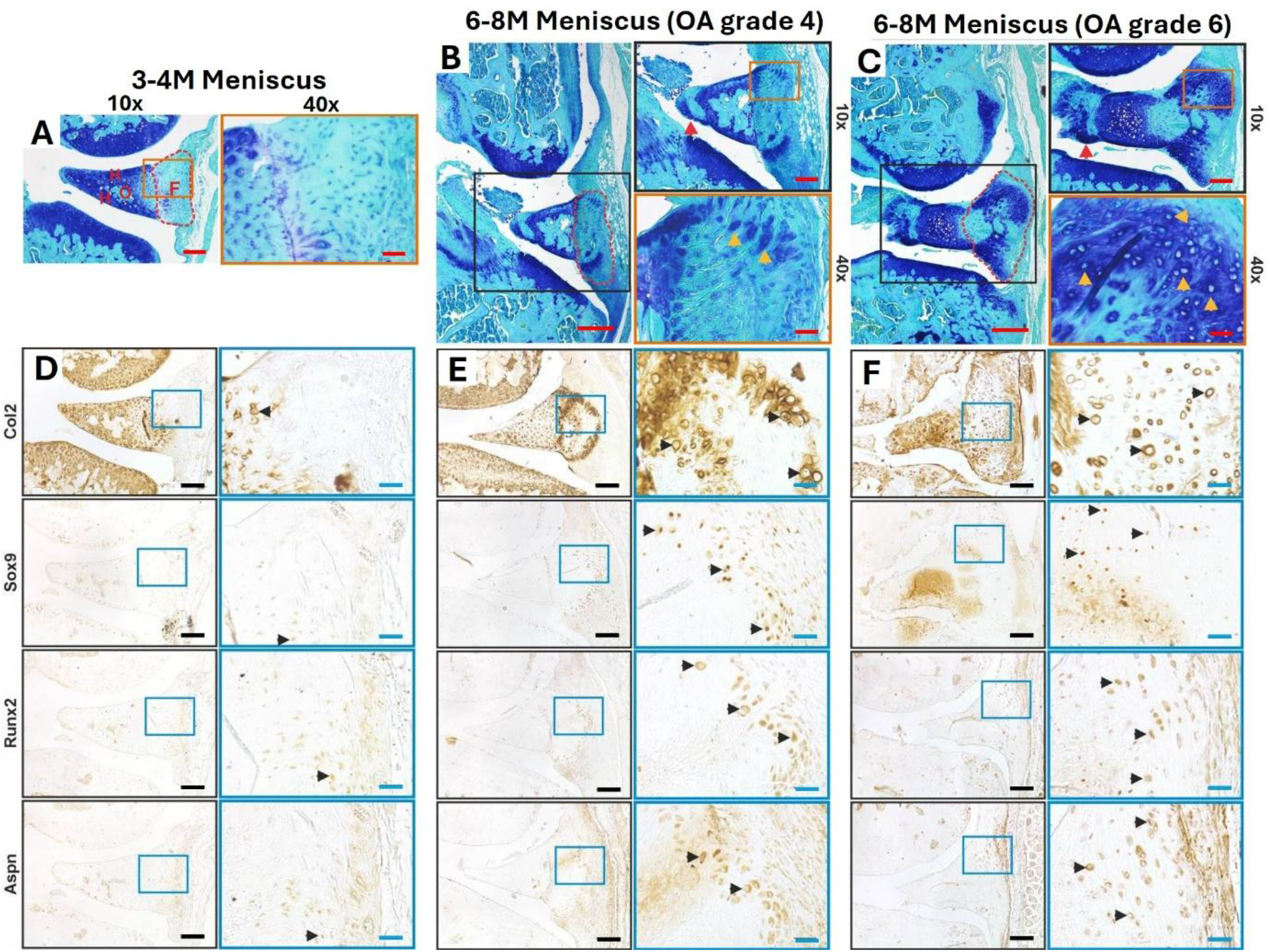
Histological toluidine blue and immuno-staining of the medial meniscus in healthy and OA knee joints of CD1 mice. Representative images: (A) healthy 3-month-old CD1 medial menisci with typical structure including including hyaline cartilage regions, ossified core and outer fibrous region (indicated as H, O and red region F, respectively) (B) OA grade 4 with expanded fibrous areas (red region) with toluidine blue staining and rounded-cell morphology (yellow arrows), as well as potential osteophyte formation at the anterior horn (red arrow). (C) OA grade 6 with enlarged fibrous regions (red region) with abundant toluidine blue staining and rounded-cell morphology (yellow arrows), and potential osteophyte formation at the anterior horn (red arrow). **(D)** COL2, SOX9, RUNX2 and ASPN immunostaining of the medial menisci of healthy 3-month-old CD1 and (E) in OA grade 4 and (F) grade6. Black arrows point to examples of positive cellular staining. Scale is 200 µm (4x), 100 µm (10x) and 25 µm (40x).

### CD1 mice at 6-8 months-old have significantly reduced fiber cross sectional area in skeletal muscles, which correlates with OA lesion severity

Although there was reduced cross sectional area (CSA) of the muscle fibers in the quadriceps (78.85±3.688 mm/MW to 72.05±1.826 mm/MW), this was not statistically significant (p=0.097; Fig.5A-C). Fiber CSA in the gastrocnemius, however, was significantly reduced at 6-8-months-old, compared to the 3-4-month-old mice (76.66±2.243 mm/Mouse Weight (MW) at 3-4M to 61.44±2.222 mm/MW at 6-8M; p<0.0001; Fig.5A-C). This reduction of CSA in the gastrocnemius, was significantly correlated with maximum OA severity (Fig.5B, p=0.0054).

**Fig. 5:**
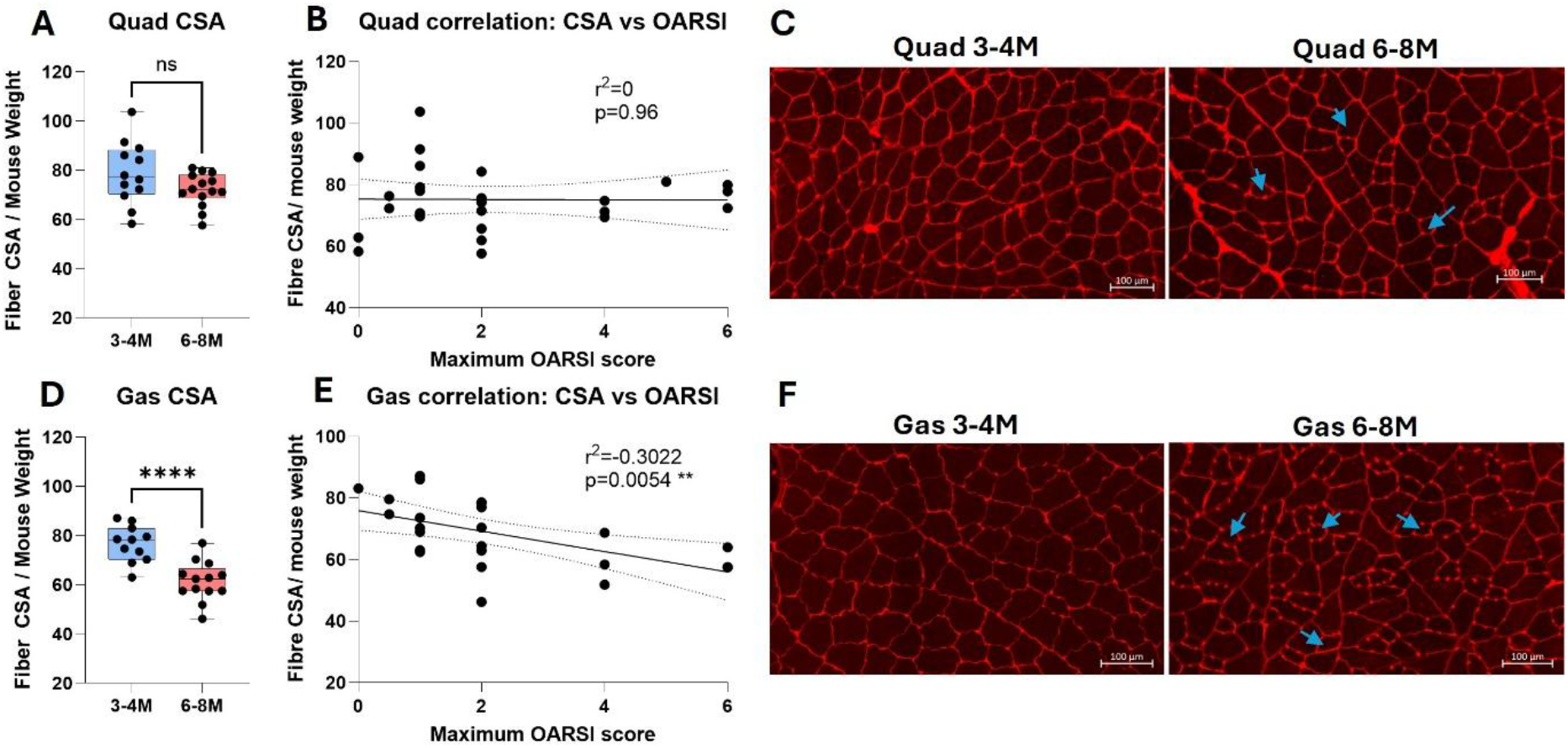
Quadriceps (Quad) and Gastrocnemius (Gas) muscle fibres cross-sectional area (CSA) in CD1 mice. **(A,D**) Fiber cross-sectional area normalized to mouse weight in the Quad (A) and Gas (D) of CD1 mice comparing 3-4 months with 6-8 months-old tissues. (B,E) Pearson’s correlation of fiber CSA (normalized to mouse weight) to the maximum OARSI score in the joint in Quad (B) and Gas (E). (C,F) representative images of cross-sections of the Quad (C) and Gas (F) in 3-4 months and 6-8 months-old CD1 mice. Blue arrows indicate smaller muscle fibers throughout the section. Student’s unpaired t-test. ns = not significant, ****p<0.0001.

## Discussion

This study describes the CD1 outbred strain of mouse as a new model of spontaneous OA. In addition to the articular cartilage loss and synovial hyperplasia that is often reported in other OA models, this study characterises novel OA associated ligament and meniscal pathological endochondral ossification, and peri-articular muscle wasting. This supports the use of CD1 male mice as a new model of spontaneous OA, with all musculoskeletal hallmarks of OA development, similar to other murine models of OA OA severity in the CD1 mice was assessed by OARSI scoring (Glasson et al., 2010) for articular cartilage lesion severity. We found significant OA development by 6 months of age, with only mild disease at 3-4 months of age. These findings point to a steep OA progression in 2 months, between 4 and 6 months of age, suggesting these timepoints as potential targets for future studies using this model to study OA development and to test potential therapies. This is comparable to the Str/ort mouse model that is the most used model of spontaneous murine OA (Poulet et al., 2012; Staines et al., 2017). Indeed, male Str/ort mice develop OA spontaneously with human-like cartilage lesions at approximately 12 to 20 weeks of age (Mason et al., 2001), primarily in the medial compartment, as well as ossification of the medial meniscus and medial collateral ligament calcification (Ramos-Mucci et al., 2020; Walton, 1977); all features that we also described in the CD1 mice in this study.

The anterior cruciate (ACL) and collateral ligaments (CLs), and medial menisci of the CD1 mice showed increases in toluidine blue staining, with OA development, suggestive of proteoglycan and glycosaminoglycan deposition (Schmitz et al., 2010). To confirm the development of pathological endochondral ossification in these soft tissues, we performed immunohistochemistry with markers of endochondral ossification (COL2, SOX9 and RUNX2), whist ASPN was chosen as a known inhibitor of chondrogenesis in cartilage and in OA (Kizawa et al., 2005) and a known ligament and tendon proteoglycan(Kharaz et al., 2016). Diseased ligaments and menisci in osteoarthritic CD1 knees showed COL2 deposition and SOX9, and RUNX2 expressing cells in areas of hyperplasia in the ACL tibial enthesis, CL enthesis, mid-ligament and regions, and the medial menisci fibrous region, markers of endochondral ossification. Similar pathologies have been described in other murine models of OA, including the Str/ort mouse (spontaneous OA), as well as injury induced OA models such as the destabilisation of the medial meniscus (DMM model) and the joint overloading model (Ramos-Mucci et al., 2020). The ligament pathologies found in our study have been linked to abnormal mechanical properties of the ACL in such models, suggesting their importance in the development and progression of OA (Ramos-Mucci et al., 2022). They resemble pathological mineralisation in human OA ligaments and entheses (Asahara et al., 2017; Benjamin and McGonagle, 2009), supporting the translational relevance of CD1 mice for studying whole-joint tissue interactions. Similarly, meniscal pathology in CD1 mice also aligns with human OA, where fibrous region expansion, proteoglycan accumulation, and osteophyte formation are common features (Fox et al., 2015).

Muscle wasting was also demonstrated from 6-8 months of age in the gastrocnemius muscles, which significantly correlated with OA severity in the joint. This finding ultimately supports the relationship between OA and muscle wasting, suggesting that induction of spontaneous OA leads to atrophy in the muscles surrounding the joint. Interestingly, the gastrocnemius experienced significantly more atrophy than the quadriceps muscles in the 6-8 month-old CD1 mice. One possible explanation for the reduced gastrocnemius fibre CSA is altered limb use associated with OA-related joint pathology. The gastrocnemius contributes substantially to locomotion in rodents, contributing to both ankle and knee mechanics during walking (Alexander et al., 2021; Baltgalvis et al., 2012). In a surgical model of OA in rats (ACL transection), the gastrocnemius showed measurable muscle loss, with about 10% reduction in gastrocnemius cross-sectional area alongside increased inflammatory signalling (Silva et al., 2018). However, locomotor activity and weight bearing were not assessed in the present study, and this possibility therefore requires further investigation The CD1 model as described herein captures the multi-tissue nature of OA, supporting the view that OA is a disease of the entire joint organ (Loeser et al., 2012). The simultaneous involvement of cartilage, synovium, ligaments, menisci, and muscles mirrors human OA pathology and provides a platform for studying cross-tissue signalling. This whole-joint phenotype is increasingly recognised as central to OA pathogenesis, where mechanical loading, inflammatory mediators and matrix-derived signals propagate between tissues to amplify degeneration (Hunter and Bierma-Zeinstra, 2019). For example, synovial inflammation can accelerate cartilage breakdown through cytokine-driven catabolism (Scanzello and Goldring, 2012), while ligament and meniscal ossification alters joint biomechanics and increases focal cartilage stress (Fox et al., 2015). Muscle wasting further contributes to aberrant joint loading and reduced shock absorption, creating a feedback loop that exacerbates structural deterioration (Shorter et al., 2019). By recapitulating these possibly interconnected pathological processes, CD1 mice offer a valuable system for dissecting how tissue-to-tissue communication drives OA progression and for identifying therapeutic targets that address OA as a coordinated organ-level disease rather than a cartilage-centric disorder.

Our study proposes the CD1 strain of mice as a novel model of spontaneous knee OA. Using the CD1 as a model offers the advantage that it is readily available around the world, usually at low cost compared to other mouse strains. In addition, CD1 mice are often used in animalfacilities as sentinels or as surrogate parents for the creation of new transgenic lines. Tissues are often left unused from these animals, but collecting and using the joints for studies of OA would provide an important reduction in the use of experimental animals, thereby supporting the 3Rs of animal experimentation. Finally, CD1 mice are an outbred strain and therefore have a more diverse genetic background compared to commonly used inbred strains, which may enhance their utility for studying osteoarthritis (Aldinger et al., 2009). Genetic heterogeneity more closely reflects the polygenic and multifactorial nature of human OA, where numerous interacting variants influence susceptibility, tissue-specific degeneration and progression rates (Reynard and Loughlin, 2012). Outbred strains have been shown to exhibit greater phenotypic variability, offering a model that captures the spectrum of OA severity and multi-tissue involvement observed clinically rather than a single, genetically constrained trajectory (Tuttle et al., 2018). This diversity also increases the likelihood of identifying gene–environment interactions, including modifiers of cartilage breakdown, synovial inflammation, ligament ossification and muscle wasting; key processes highlighted in our CD1 cohort. Furthermore, outbred mice may improve translational relevance for therapeutic testing, as treatment responses can be evaluated across a broader biological background, better approximating genetically diverse human populations. Collectively, the genetic diversity of CD1 mice strengthens their value as a spontaneous OA model capable of revealing mechanisms that may be obscured in highly inbred strains.

## Conclusions

Together, these findings establish the CD1 outbred mouse as a robust and biologically relevant model of spontaneous osteoarthritis. CD1 mice develop progressive, multi-tissue joint pathology—including articular cartilage degeneration, synovial inflammation, ligament and meniscal endochondral ossification, and peri-articular muscle wasting—that closely mirrors the complex organ-level nature of human OA. Their genetic diversity, global availability, and alignment with the 3Rs further enhance their practical and translational value. By capturing the coordinated deterioration of multiple joint tissues and reflecting the polygenic variability of human disease, the CD1 model provides a powerful platform for investigating cross-tissue mechanisms, identifying molecular drivers of OA progression, and evaluating therapeutic strategies aimed at restoring whole-joint health.

## Acknowledgements

The Authors acknowledge use of the University of Liverpool Histology Shared Research Facility (RRID:SCR_026606).

## Author contributions statement

Conceptualization: B.P; Methodology: B.P., K.G.W., E.C., T.D., E.S., L.R.M.; Validation: all authors; Formal analysis: E.S., L.R.M., C.K., T.D., Y.Wen., Y.Wang.; Writing - original draft: B.P., E.S., L.R.M.; Writing - review & editing: all authors; Visualization: B.P., E.S., L.R.M., T.D.; Supervision: B.P., K.G.W., E.C.; Funding acquisition: B.P., K.G.W., E.C., Y.Wen.

## References

A. Ratneswaran, B. T. B.A. Russell, F. Beier,. (2018). Strain and model specific differences in mouse models of osteoarthritis. Osteoarthritis and Cartilage 26, S71.

Aldinger, K. A., Sokoloff, G., Rosenberg, D. M., Palmer, A. A. and Millen, K. J. (2009). Genetic variation and population substructure in outbred CD-1 mice: implications for genome-wide association studies. PLoS One 4, e4729.

Alexander, P., Vsevolod, L., Natalia, M. and Pavel, M. (2021). Effect of hindlimb unloading on recruitment of gastrocnemius medialis muscle during treadmill locomotion in rats. Exp Brain Res 239, 2793–2801.

Asahara, H., Inui, M. and Lotz, M. K. (2017). Tendons and Ligaments: Connecting Developmental Biology to Musculoskeletal Disease Pathogenesis. J Bone Miner Res 32, 1773–1782.

Baltgalvis, K. A., Call, J. A., Cochrane, G. D., Laker, R. C., Yan, Z. and Lowe, D. A. (2012). Exercise training improves plantar flexor muscle function in mdx mice. Med Sci Sports Exerc 44, 1671–9.

Benjamin, M. and McGonagle, D. (2009). Entheses: tendon and ligament attachment sites. Scand J Med Sci Sports 19, 520–7.

Fox, A. J., Wanivenhaus, F., Burge, A. J., Warren, R. F. and Rodeo, S. A. (2015). The human meniscus: a review of anatomy, function, injury, and advances in treatment. Clin Anat 28, 269–87.

Glasson, S. S., Chambers, M. G., Van Den Berg, W. B. and Little, C. B. (2010). The OARSI histopathology initiative -recommendations for histological assessments of osteoarthritis in the mouse. Osteoarthritis Cartilage 18 Suppl 3, S17–23.

Hunter, D. J. and Bierma-Zeinstra, S. (2019). Osteoarthritis. Lancet 393, 1745–1759.

Johnson, V. L. and Hunter, D. J. (2014). The epidemiology of osteoarthritis. Best Pract Res Clin Rheumatol 28, 5–15.

Kharaz, Y. A., Tew, S. R., Peffers, M., Canty-Laird, E. G. and Comerford, E. (2016). Proteomic differences between native and tissue-engineered tendon and ligament. Proteomics 16, 1547–56.

Kizawa, H., Kou, I., Iida, A., Sudo, A., Miyamoto, Y., Fukuda, A., Mabuchi, A., Kotani, A., Kawakami, A., Yamamoto, S. et al. (2005). An aspartic acid repeat polymorphism in asporin inhibits chondrogenesis and increases susceptibility to osteoarthritis. Nat Genet 37, 138–44.

Kuyinu, E. L., Narayanan, G., Nair, L. S. and Laurencin, C. T. (2016). Animal models of osteoarthritis: classification, update, and measurement of outcomes. J Orthop Surg Res 11, 19.

Loeser, R. F. (2010). Age-related changes in the musculoskeletal system and the development of osteoarthritis. Clin Geriatr Med 26, 371–86.

Loeser, R. F., Goldring, S. R., Scanzello, C. R. and Goldring, M. B. (2012). Osteoarthritis: a disease of the joint as an organ. Arthritis Rheum 64, 1697–707.

Man, G. S. and Mologhianu, G. (2014). Osteoarthritis pathogenesis -a complex process that involves the entire joint. J Med Life 7, 37–41.

Mason, R. M., Chambers, M. G., Flannelly, J., Gaffen, J. D., Dudhia, J. and Bayliss, M. T. (2001). The STR/ort mouse and its use as a model of osteoarthritis. Osteoarthritis Cartilage 9, 85–91.

Obeidat, A. M., Kim, S. Y., Burt, K. G., Hu, B., Li, J., Ishihara, S., Xiao, R., Miller, R. E., Little, C., Malfait, A. M. et al. (2024). A standardized approach to evaluation and reporting of synovial histopathology in two surgically induced murine models of osteoarthritis. Osteoarthritis Cartilage 32, 1273–1282.

Poulet, B., Hamilton, R. W., Shefelbine, S. and Pitsillides, A. A. (2011). Characterizing a novel and adjustable noninvasive murine joint loading model. Arthritis Rheum 63, 137–47.

Poulet, B., Ulici, V., Stone, T. C., Pead, M., Gburcik, V., Constantinou, E., Palmer, D. B., Beier, F., Timmons, J. A. and Pitsillides, A. A. (2012). Time-series transcriptional profiling yields new perspectives on susceptibility to murine osteoarthritis. Arthritis Rheum 64, 3256–66.

Ramos-Mucci, L., Elsheikh, A., Keenan, C., Eliasy, A., D’Aout, K., Bou-Gharios, G., Comerford, E. and Poulet, B. (2022). The anterior cruciate ligament in murine post-traumatic osteoarthritis: markers and mechanics. Arthritis Res Ther 24, 128.

Ramos-Mucci, L., Javaheri, B., van ‘t Hof, R., Bou-Gharios, G., Pitsillides, A. A., Comerford, E. and Poulet, B. (2020). Meniscal and ligament modifications in spontaneous and post-traumatic mouse models of osteoarthritis. Arthritis Res Ther 22, 171.

Reynard, L. N. and Loughlin, J. (2012). Genetics and epigenetics of osteoarthritis. Maturitas 71, 200–4

Scanzello, C. R. and Goldring, S. R. (2012). The role of synovitis in osteoarthritis pathogenesis. Bone 51, 249–57.

Schmitz, N., Laverty, S., Kraus, V. B. and Aigner, T. (2010). Basic methods in histopathology of joint tissues. Osteoarthritis Cartilage 18 Suppl 3, S113–6.

Shorter, E., Sannicandro, A. J., Poulet, B. and Goljanek-Whysall, K. (2019). Skeletal Muscle Wasting and Its Relationship With Osteoarthritis: a Mini-Review of Mechanisms and Current Interventions. Curr Rheumatol Rep 21, 40.

Silva, J. M. S., Alabarse, P. V. G., Teixeira, V. O. N., Freitas, E. C., de Oliveira, F. H., Chakr, R. and Xavier, R. M. (2018). Muscle wasting in osteoarthritis model induced by anterior cruciate ligament transection. PLoS One 13, e0196682.

Staines, K. A., Poulet, B., Wentworth, D. N. and Pitsillides, A. A. (2017). The STR/ort mouse model of spontaneous osteoarthritis -an update. Osteoarthritis Cartilage 25, 802–808.

Terry, R. L. and Wells, D. J. (2016). Histopathological Evaluation of Skeletal Muscle with Specific Reference to Mouse Models of Muscular Dystrophy. Curr Protoc Mouse Biol 6, 343–363.

Tuttle, A. H., Philip, V. M., Chesler, E. J. and Mogil, J. S. (2018). Comparing phenotypic variation between inbred and outbred mice. Nat Methods 15, 994–996.

Walton, M. (1977). Degenerative joint disease in the mouse knee; radiological and morphological observations. J Pathol 123, 97–107.

Wen, Y., Murach, K. A., Vechetti, I. J., Jr., Fry, C. S., Vickery, C., Peterson, C. A., McCarthy, J.J. and Campbell, K. S. (2018). MyoVision: software for automated high-content analysis of skeletal muscle immunohistochemistry. J Appl Physiol (1985) 124, 40–51.

