## Supplementary figures for "Pathological musculoskeletal characterisation of a new model of spontaneous OA in CD1 mice"

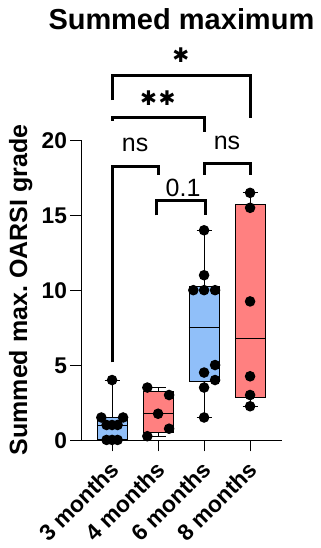

**Supplementary Figure 1**: Summed maximum OARSI scores for each timepoint separately, showing similar average scores between 3 and 4 months and between 6 and 8 months-old mice. These groups are subsequently grouped to increase power for young healthy 3-4 month-old mice and older osteoarthritic 6-8 months-old.

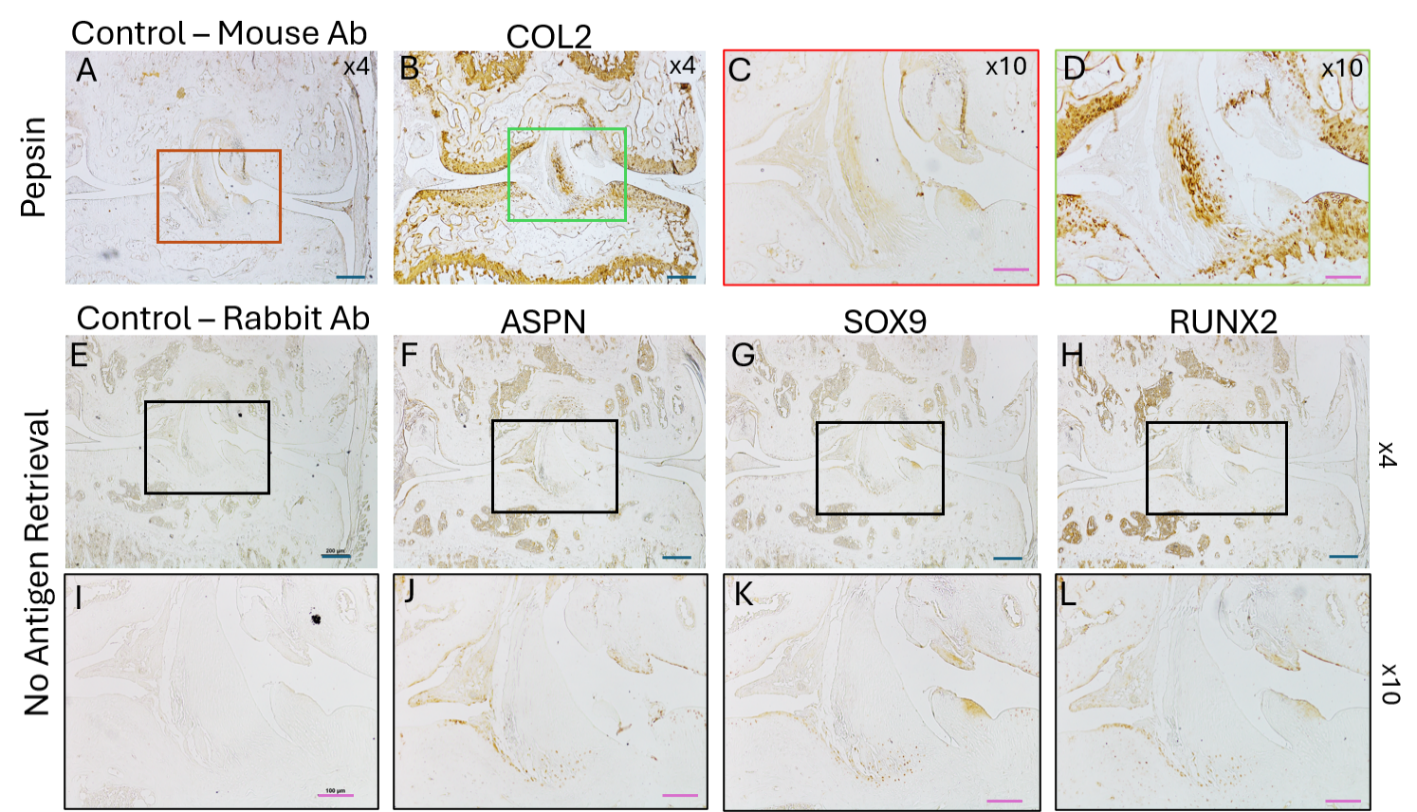

**Supplementary Figure 2:** Representative images of negative controls for COL2 staining (mouse IgG, Pepsin as antigen retrieval) and for ASPN, SOX9 and RUNX2 (rabbit IgG, no antigen retrieval). Images shown at x4 of the whole knee joint, and x10 magnification of the anterior cruciate ligament area. Scale bars: blue = 200µm; pink = 100µm. All images in this figure are from the same joint and adjacent sections, and shows very limited unspecific staining with both mouse and rabbit primary antibodies.
